# Chronic Intermittent Ethanol Exposure Produces Physiological Tolerance and Distinct Behavioral Alterations Across Abstinence in Female Mice

**DOI:** 10.64898/2026.09.29.755351

**Authors:** Maya R. Eberle-Redding, Ann M. Walker, Shivani P. Vaidya, Jobe L. Ritchie, Lisa R. Taxier, Baylee R. Materia, Leon G. Coleman, Ryan P. Vetreno, Thomas L. Kash, Brianna E. George

## Abstract

Chronic ethanol consumption produces long-lasting effects on the brain that contribute to persistent behavioral changes. This has been modeled in animals using the chronic intermittent ethanol (CIE) paradigm, which induces escalation of consumption and alterations in affective and cognitive behavior. While this model has been well characterized in male subjects, the effects of CIE in females and long-term effects that persist beyond the acute withdrawal remain relatively underexplored. Here, female C57BL/6J mice were exposed to four weeks of chronic intermittent ethanol (CIE) vapor to induce dependence. Following CIE, mice were evaluated for physiological tolerance to ethanol and alterations in anxiety-like behaviors and memory processing. Using the ethanol responsivity battery (ERB) assay to assess physiological tolerance, we found blunted responsiveness to the intoxicating effects of ethanol in the CIE-exposed females, indicating heightened ethanol tolerance that persists beyond the acute withdrawal phase. A cohort underwent behavioral testing following CIE to characterize changes in anxiety-like behavior, exploration, and memory across abstinence. In early abstinence (<5 days post-CIE), CIE-exposed females showed increased exploratory behavior compared with Air controls. After one week of abstinence, CIE-exposed females displayed impaired object recognition memory. CIE-exposed females also showed increased anxiety-like behavior at protracted abstinence time points, exhibiting hyponeophagia in the novelty-suppressed feeding test at 2 weeks post-CIE. In addition, following cued fear conditioning, we found reduced extinction recall at 4 weeks post-CIE. Together, this work indicates that chronic alcohol exposure leads to increased alcohol tolerance and distinct, persistent alterations in memory and anxiety-like behavior in female mice.

## Introduction

Alcohol use disorder (AUD) is a highly prevalent disorder within the United States, with approximately 25.7 million people ages 12 and older having an AUD diagnosis in 2025^1^. Although men have higher incidence rates of AUD^1^, between 2002 and 2013, there was an 83.7% increase in the number of women with AUD^2^. To further highlight sex differences in the context of AUD, sex has a profound effect on withdrawal symptomology. During withdrawal, which is a defining feature of AUD^3^, people can experience sleep disturbances, cognitive impairments, cravings for alcohol, and hyperkatifeia, amidst other symptoms of negative affect^3,4^. These experiences may drive a person to reinstate their consumption of ethanol^5^, lending to a cyclic nature of intoxication and withdrawal^4^. In the past, women have been found to have increased cravings for alcohol^6,7^, as well as higher percentages of comorbid alcohol dependence and anxiety/affective disorders^8,9^. However, other studies have shown greater reports of alcohol withdrawal-related anxiety^10^ and higher withdrawal severity scores in men^11^. These differences in findings can be attributed to the complex nature of AUD and withdrawal, as well as the historical underrepresentation of women and inclusion of both sexes within alcohol research^12,13^. Given the current rise in the number of women diagnosed with AUD^14^, understanding the behavioral manifestations of AUD in females, especially during the critical withdrawal period that may increase relapse vulnerability, is warranted.

To study AUD, many animal models have been developed to examine distinct aspects of the condition. The chronic intermittent ethanol (CIE) procedure is a model used to assess alcohol dependence. In this procedure, rodents are exposed to a series of ethanol intoxication sessions, using non-contingent delivery of ethanol vapor, followed by intermittent periods of withdrawal^15^. Prior studies have looked at the behavioral impact of abstinence from ethanol following CIE exposure in rodents, with a focus primarily in males. One key finding that points towards the mechanistic validity of the CIE model is that it drives increased voluntary ethanol consumption^16,17^. It has been hypothesized that this is due to heightened negative affective states, which have been a major focus of behavioral analysis following CIE. During the withdrawal period, male rats exposed to 10 days of CIE vapor show an increase in anxiety-like behaviors in the light-dark box^18^and the elevated plus maze assays during abstinence^19^. Similarly, after CIE, male C57BL/6J mice displayed increased anxiety-like behaviors during ethanol abstinence in the novelty suppressed feeding^20–22^ and marble burying assays^20,21,23–25^. Interestingly, there are mixed results when longitudinally assessing the impact of protracted CIE abstinence on behavior, with one paper reporting a decrease of anxiety-like behavior^26^, while another shows persistence of this anxiety-like state across the acute withdrawal and protracted abstinence periods^27^. In addition to impacting anxiety-like behaviors, ethanol abstinence was found to cause changes to spatial^28^ and non-spatial memory^29^ in rats. Importantly, many of these studies did not use female rodents as subjects or found little to no impact of ethanol abstinence on behavior in females^19,21^, leaving a substantial gap in the literature regarding the behavioral changes in females caused by ethanol abstinence.

To fill this knowledge gap, the goals of the present study were to evaluate if CIE increases alcohol tolerance, and drives aberrant behavioral changes of CIE-exposed female mice across time. To accomplish the first goal, alcohol tolerance was analyzed following the CIE paradigm using the ethanol responsivity battery (ERB)^30–32^, which is a previously established paradigm that assesses intoxication through a variety of tests. The second goal was then addressed by using a four-week CIE paradigm followed by a battery of behavioral assays. Behavioral testing was performed for five weeks to examine changes in anxiety-like states, exploration, and memory across time. By characterizing these behaviors across weeks, this paper aims to identify the changes resulting from chronic ethanol exposure and abstinence. The results of this study may provide insight into the growing incidence of women with AUD by specifically highlighting abstinence-induced negative affective states that may cause a reinstatement of alcohol use.

## Methods

### Animals

Female (*n* = 33) C57BL/6J mice were obtained from the Jackson Laboratory (Bar Harbor, ME, USA) at seven weeks old and were group-housed prior to the start of the experiment. Mice were housed in static cages and placed in the vapor chambers for acclimation one week before the chronic intermittent ethanol (CIE) paradigm began. Throughout the experiment, mice were maintained on a 12-hour reverse light-dark cycle (lights off at 7AM) and were allowed *ad libitum* access to food (Prolab Isopro RMH 3000, LabDiet) and water. All procedures were performed as instructed by the National Institute of Health’s Guide for the Care and Use of Laboratory Animals and were approved by the Institutional Animal Care and Use Committee at the University of North Carolina at Chapel Hill.

### Chronic Intermittent Ethanol

The chronic intermittent ethanol (CIE) paradigm was used to administer ethanol to mice via inhalation as previously described^15,33,34^. In this experiment, mice were randomly assigned to a condition and placed in either air (AIR, *n* = 16) or ethanol vapor chambers (CIE, *n* = 17) for 16 hours of air or ethanol vapor exposure, on a four-days on, three-days off schedule. This was repeated four times, for a total of four CIE cycles. Prior to exposure, each mouse received an intraperitoneal (i.p.) injection of either a 1 mM pyrazole and saline solution (AIR), or a 1 mM pyrazole, saline, and 95% ethanol (1.6 g/kg) solution (CIE). Pyrazole solutions were kept at 4°C and brought to room temperature ∼1 hour before injections took place. After vapor exposure, all mice were kept in room air for 8 hours. On the first week of CIE, the vapor rate was kept low to reduce the chance of adverse outcomes. However, for the following weeks, the flow rate was adjusted to maintain vapor levels that produced average blood ethanol concentrations (BECs) above 0.08%, the defined binge alcohol exposure percentage per the National Institute on Alcohol Abuse and Alcoholism^35^. At the end of each week, ∼10 µL of tail blood was collected from CIE mice only, and BECs were quantified using the Analox AM1 Alcohol Analyzer (Analox Instruments Ltd., UK). Samples contaminated during the weekly collection were excluded from group values. Once the paradigm was complete, 6 AIR-exposed and 7 CIE-exposed mice were selected to carry out ethanol responsivity testing (Experiment 1). One CIE mouse was excluded from this testing due to adverse outcomes. The remaining mice (*n* = 20) were tested in a battery of behavioral assays to assess anxiety-like states, exploration, and memory performance post-CIE (Experiment 2).

#### Experiment 1: Ethanol Response Battery

The ethanol response battery (ERB) was carried out on 6 AIR-exposed and 6 CIE-exposed females beginning seven days after the conclusion of CIE treatment, and was performed as previously described^30–32^. After a baseline assessment including body weight measurements, mice were given four consecutive intraperitoneal (i.p.) injections of ethanol (i.e. 0.5, 0.5, 1.0, 1.0 g/kg), which produced cumulative ethanol doses of approximately 0.5, 1.0, 2.0, and 3.0 g/kg. This dosing regimen permits evaluation of ethanol responsivity during the rising phase of the BECs, across a broad range of ethanol doses. The ERB assessment was carried out approximately 15 minutes after each ethanol dose, and included the behavioral intoxication rating scale, body temperature assessment, submandibular blood collection, and tilting plane motor coordination assay. The loss of righting reflex (LORR) assay was only carried out after the final ethanol dose.

### Behavioral Intoxication Rating Scale

The behavioral intoxication rating scale was utilized as previously described^30^, where mice were rated separately by two researchers on a 6-point scale of no intoxication to extreme intoxication. The breakdown of the scale is as follows: (1) no sign of intoxication; (2) hypoactivity; (3) slight intoxication (ataxia; slight motor impairment); (4) moderate intoxication (obvious motor impairment; dragging abdomen); (5) high intoxication (dragging abdomen; LORR); (6) extreme intoxication (LORR; loss of eye blink response). Scores were averaged between the two researchers and recorded.

### Body Temperature Assessment

At baseline and following each behavioral intoxication rating, body temperature was assessed using a Physitemp Thermalert clinical monitoring thermometer (Physitemp Instruments, Clifton, NJ). To obtain an accurate and stable temperature reading, animals were restrained for 2 minutes in a DecapiCone (ThermoFisher Scientific, Austin, TX) while an electric thermometer probe was inserted approximately 3 mm into the rectum of each mouse and left in place for ≥ 45 seconds. Changes in body temperature were calculated by subtracting body temperature following each cumulative ethanol dose from the baseline temperature. Ambient room temperature was also monitored and averaged 23°C.

### Blood Collection

Submandibular blood was collected from mice at baseline, as well as after body temperature assessment to determine their BECs as mice received cumulative ethanol doses. Serum was analyzed using the Analox GM7 Analyzer (Analox Instruments Ltd., UK).

### Tilting Plane

Mice were administered the tilting plane assay at baseline and after each ethanol dose during ERB testing. The tilting plane apparatus consisted of a clear Plexiglas box (18 cm × 7.5 cm × 9.5 cm), which was attached with a hinge to a framed glass panel floor. An additional hinge was used to tilt the box, and the angle was measured by a digital angle gauge. At the beginning of the test, the mouse was placed inside the box facing away from the tilting hinge. The panel floor lifted slowly until the mouse began to slide down the floor of the apparatus, at which point the angle was measured and recorded. This was repeated three times per session. The three trials within each session were averaged and the difference in angle of slide as a consequence of cumulative ethanol dosing was calculated by subtracting angle of slide from the average baseline angle of slide from each ethanol dose angle of slide.

### Loss of Righting Reflex

After mice were given the final dose of ethanol, and had completed the tilting plane assay, loss of righting reflex (LORR) was assessed. Mice were placed on their back in a V-shaped trough, and were scored on their ability to right themselves. LORR was defined as the inability of the mouse to right itself onto all four paws within 60 seconds.

#### Experiment 2: Post-CIE Behavioral Testing

A separate group of 10 AIR-exposed and 10 CIE-exposed mice were administered five weeks of behavioral testing following the end of CIE treatment. Testing began three days after CIE- or AIR-exposure and continued until post-CIE Day 27.

### Elevated Zero Maze

The elevated zero maze (EZM) assay was conducted on Day 3 of the post-CIE period, on an elevated circle-shaped maze (Med Associates, Product Number ENV-561A-BL) with a diameter of 61 cm. Two quadrants on opposing sides of the zero maze are enclosed by blue walls standing ∼8 cm high (closed zones), while the other two quadrants are left open (open zones). The maze was illuminated by four LED strips mounted on the ceiling, to produce a lux of ∼32 in the open quadrants, and ∼28 in the closed. Mice were placed in an open zone, with their face oriented toward the enclosed area, and were allowed to explore the maze for 10 minutes. The maze was cleaned with a 70% ethanol solution between mice. All behavior was recorded and tracked using Noldus EthoVision software (Noldus Information Technology BV, Wageningen, Netherlands).

### Open Field Test

The open field test (OFT) assay was conducted in a white Plexiglas box with an open top, measuring 50 cm by 50 cm, 5 days after CIE. The box was illuminated by two LED light strips positioned above the box, to reach a lux of ∼30 in the center. Each mouse was placed in the bottom corner of the box, with their head oriented toward the corner, and their behavior was recorded for 30 minutes. A 70% ethanol solution was used to clean the box between mice. All behavior was recorded and tracked using Noldus EthoVision software (Noldus Information Technology BV, Wageningen, Netherlands). Behavior was then analyzed using DeepLabCut for behavioral pose estimation and processed using ABEL^36^, to quantify exploration, rearing, and grooming behaviors.

### Novel Object Recognition

The novel object recognition (NOR) assay was carried out on Days 6 and 7 post-CIE. On both days, mice were recorded in a white Plexiglas box with an open top, measuring 50 cm by 50 cm. The box was illuminated from above by two LED light strips to obtain a lux of ∼30 in the center. As all mice had previous experience with the testing box during the OFT, we did not run a separate acclimation session. At the start of the recording, each mouse was placed in the bottom corner of the box, with their head facing the corner. Between mice, a 70% ethanol solution was used to clean the box and objects. All behavior was recorded and tracked using Noldus EthoVision software (Noldus Information Technology BV, Wageningen, Netherlands). On the first day of this assay, mice were given 10 minutes to explore either two locks or two pipes, which were placed on opposite corners of the box, with the center of each object located approximately 14 cm from the sides of the box (training day). On the second day, mice were permitted 10 minutes of free exploration and were tested on their ability to recognize a novel object by switching out one of the locks with a pipe, or vice versa (testing day).

### Light-Dark Box

The light-dark box (LDB) assay was performed 12 days after CIE in a ∼29 cm by ∼29 cm box consisting of an illuminated “light” side, measuring ∼175 lux in the center, and an enclosed “dark” side. Mice were placed in the corner of the light side, with their heads facing the corner, and were allowed to roam the box for 10 minutes. Both sides of the box were wiped with a 70% ethanol solution between mice.

### Novelty Suppressed Feeding

The novelty suppressed feeding (NSF) assay was carried out 14 days after the conclusion of the CIE paradigm. One day prior to the behavioral assay, mice were given a small amount of rodent diet with 60 kcal% fat (Research Diets, D12492). Four hours before the start of the test, all rodent diet and regular home cage food were removed from the home cage, while water remained accessible. Testing took place in a 50 cm by 50 cm white open-topped Plexiglas box, illuminated from above to reach a lux of ∼400. At the beginning of each test, a new pellet of rodent diet was taped to the floor in the center of the box (25 cm from each wall). Each mouse was placed in the bottom corner of the box, with their head oriented toward the food pellet, and allowed to explore the box and pellet for 10 minutes. The box was cleaned with a 70% ethanol solution between each mouse. All behavior was recorded and tracked using Noldus EthoVision software (Noldus Information Technology BV, Wageningen, Netherlands).

### Acoustic Startle

The acoustic startle assay was performed on Days 17 and 18 post-CIE, using the SR-LAB Startle Response System by San Diego Instruments (San Diego, CA). During the assay, each mouse was semi-restrained in a Plexiglas tube located inside of a sound-attenuating chamber. After a five-minute acclimation period, tones of varying intensities were played throughout the chamber and startle responses were recorded. The tones were presented in a pseudo-random order: two startle tones at 105 dB and 120 dB, a blank tone (background dB), and three pre-pulse tones at 75 dB, 80 dB, and 85 dB, with a random ITI of 30 to 60 seconds between tones. Pre-pulses occurred for 20 mS, followed by an 80-mS pause, and then a 40-mS startle tone. Each mouse received five blocks of nine trials of tones, one of each intensity, for a total of 45 tone presentations per mouse.

### Elevated Plus Maze

The elevated plus maze (EPM) assay was conducted on post-CIE Day 21, on an elevated plus-shaped maze (Med Associates, Product Number ENV-560A) measuring 76 cm by 76 cm. Two opposing arms of the plus maze were enclosed by 21-cm high walls, while the other two arms were left open. The maze was illuminated by two LED strips mounted on the ceiling, to produce a lux of ∼33 in the open arms and ∼20 in the closed. Mice were placed in an open arm, with their face oriented toward the center of the maze, and were allowed to explore for 10 minutes. The maze was cleaned with a 70% ethanol solution between mice. All behavior was recorded and tracked using Noldus EthoVision software (Noldus Information Technology BV, Wageningen, Netherlands).

### Cued Fear Conditioning

The cued fear assay was carried out on post-CIE Days 25 through 27. On the first day (acquisition day), mice were placed in an illuminated (∼80 lux) 28 cm by 24 cm modular test chamber (Med Associates, Product Number ENV-008) with a metal-rod floor (context A). During the acquisition trial, after a three-minute acclimation period, mice were subject to a 20-second tone, immediately followed by a two-second foot shock (0.6 mA). In total, each mouse received five tone-shock pairings with a 90-second break between each pair. Context A was cleaned with a 1% vanilla solution (19.5% ethanol, 79.5% water) between mice. On day two (extinction day) and day three (extinction recall day), mice were placed in a new modular test chamber (Med Associates, Product Number ENV-008) with a white Plexiglas floor (context B). On both days, after three minutes of acclimation, a 20-second tone was played 20 times, with a 60-second break between each tone presentation. Between mice, context B was cleaned with a 0.5% acetic acid in ddH_2_O solution. On all three days, behavior was recorded and tracked using Noldus EthoVision software (Noldus Information Technology BV, Wageningen, Netherlands). Freezing was evaluated using ABEL and defined as absence of movement except for respiration, and had a minimum bout duration of 1000ms

### Statistical Analyses

For both experiments, all statistical analyses were performed using GraphPad Prism 10 (GraphPad Prism Software, Boston, MA, USA). Data were analyzed using repeated measures analysis of variance (RM ANOVA) with group (CIE, AIR) as the between-subjects factor and time as the within-subjects factor, or t-tests, where appropriate. Significant main effects were followed up using Bonferroni’s *post hoc* test. To analyze the LORR data in Experiment 1, a Pearson Chi Square test was used. When values were excluded from the data, mixed effects analyses were used. Data are reported as mean ± SEM. Significance was set at *p* = 0.05.

## Results

### The CIE paradigm is sufficient to produce binge BECs

For four weeks, 17 female mice were exposed to ethanol vapor through the CIE procedure. On the first week, the rate of ethanol flow was kept low to produce blood ethanol concentrations (BECs) that would prevent any adverse outcomes. For Experiment 1 (**Fig. 2A**), this resulted in a significant difference in BECs between weeks (One-Way ANOVA, *F*_(1.351,8.105)_ = 13.78, *p* = 0.004), which Bonferroni’s *post hoc* test revealed to be a significant increase in BECs between the first week and week two (*p* = 0.0321), week three (*p* = 0.0002), and week four (*p* = 0.0009). For Experiment 2 (**Fig. 2B**) a One-Way ANOVA also revealed a significant difference in BECs across weeks (*F*_(1.482,12.35)_ = 10.81, *p* = 0.0033), but only between the first week and third week (Bonferroni’s *post hoc*, *p* = 0.0007). There were no significant differences in BECs between weeks two through four (2 vs. 3, *p* = 0.7226, 2 vs. 4, *p* > 0.9999, 3 vs. 4, *p* = 0.2224). Thus, the CIE paradigm was sufficient to produce BECs that remained at or above 200 mg/dL across subsequent weeks.

**Figure 1.**
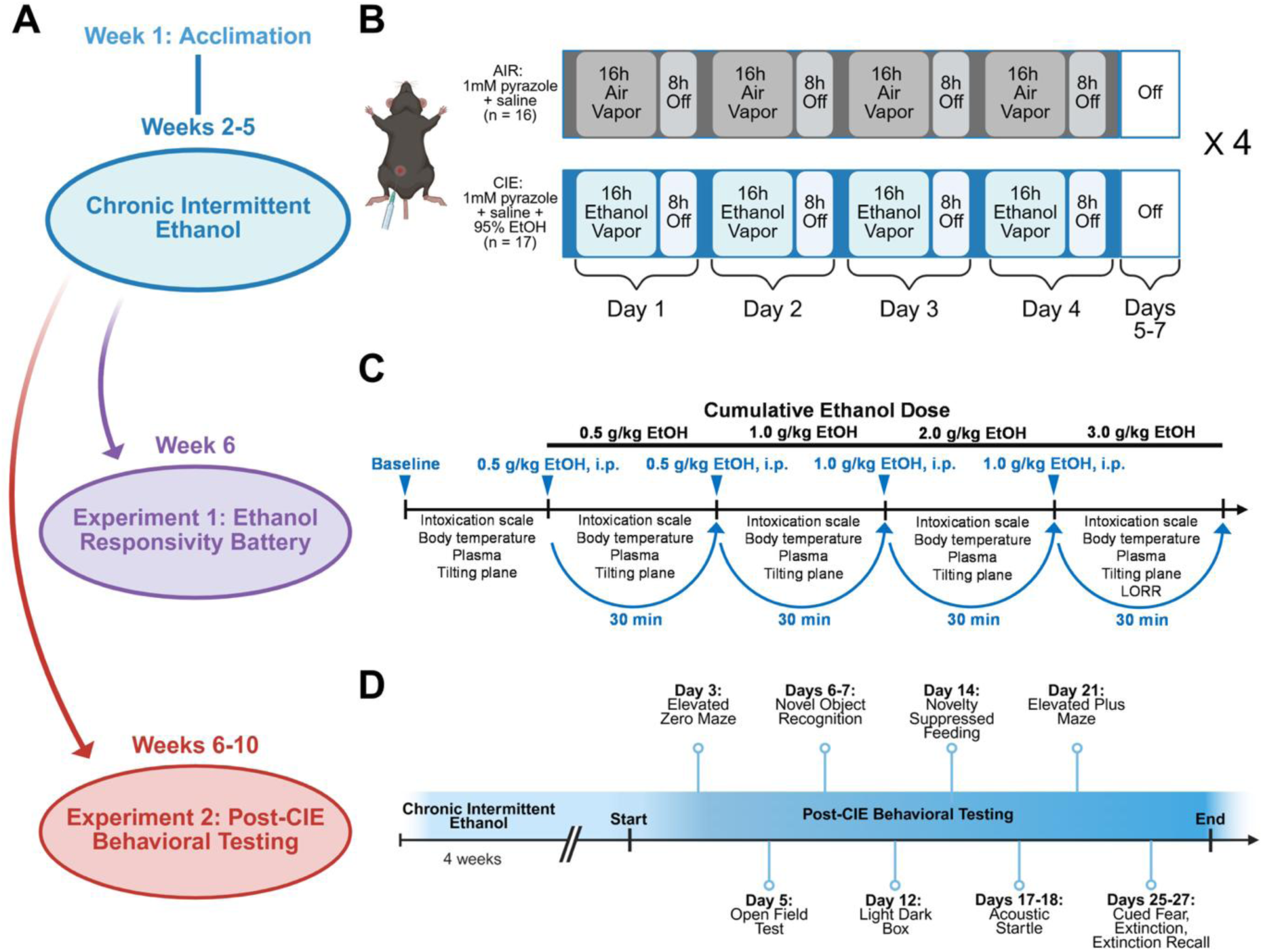
Timeline of chronic intermittent ethanol paradigm and experiments **(A)** Overall experiment timeline. 33 female mice were acclimated to living in the vapor chambers for one week before undergoing **(B)** four weeks of the chronic intermittent ethanol (CIE) paradigm, where they received either air (AIR, *n* = 16) or ethanol (CIE, *n* = 17) vapor. **(C)** Outline of the ethanol responsivity battery (Experiment 1), run on 6 AIR-exposed and 6 CIE-exposed mice one week after completing the CIE paradigm. **(D)** Timeline of the behavioral assays run post-CIE (Experiment 2). Only the mice not tested on the ethanol responsivity battery (AIR, *n* = 10, CIE, *n* = 10) were used for Experiment 2.

**Figure 2.**
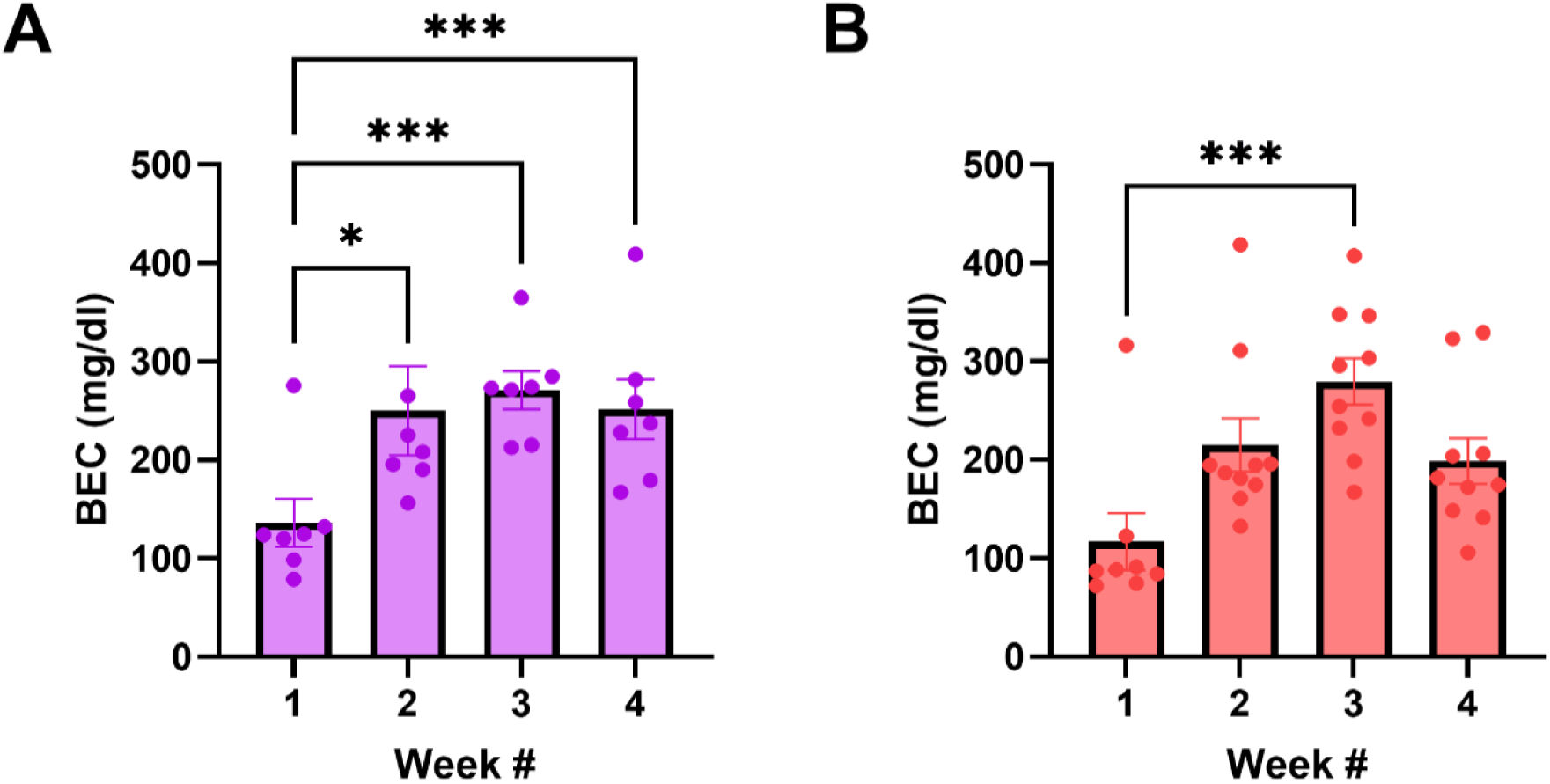
Blood ethanol concentrations for CIE-exposed mice **(A)** Blood ethanol concentrations (BECs) for the 7 CIE-exposed mice from Experiment 1. **(B)** BEC data for 10 CIE-exposed mice from Experiment 2. Data are represented as mean ± SEM.

### Ethanol Responsivity Battery: CIE-exposed mice display an increased ethanol tolerance one week after vapor exposure

The impact of four weeks of CIE on ethanol tolerance was assessed through the ethanol responsivity battery (ERB). To control for group differences, body weights were measured and analyzed prior to the start of this assay, and no significant differences were found (**Fig. 3A**, unpaired t-test, *t*(10) = 1.151, *p* = 0.2765). The ERB, completed one week after the conclusion of the CIE paradigm, revealed a significant effect of group (**Fig. 3B**, Two-Way RM ANOVA, *F*_(1,10)_ = 11.60, *p* = 0.0067), and an interaction between group and cumulative ethanol dose (Two-Way RM ANOVA, *F*_(3,30)_ = 5, *p* = 0.0063) on the intoxication ratings of mice. More specifically, Bonferroni’s *post hoc* test showed that AIR mice had significantly higher intoxication scores than the CIE mice with 1.0 (*p* = 0.0179), 2.0 (*p* = 0.0179), and 3.0 g/kg (*p* = 0.0161) of ethanol on board. After intoxication scores were recorded, change in body temperature from baseline was assessed (**Fig. 3C**). A Two-Way RM ANOVA revealed a hypothermic effect of ethanol that differed between groups (*F*_(1,10)_ = 5.956, *p* = 0.0348), where AIR mice had a significantly greater change in body temperature than CIE mice at a cumulative ethanol dose of 1.0 g/kg (Bonferroni’s *post hoc* test, *p* = 0.0272), which became a trend at 2.0 (*p* = 0.0756) and 3.0 g/kg (*p* = 0.0785) doses. A series of Two-Way RM ANOVAs uncovered no significant group differences for rising BECs (**Fig. 3D**, *F*_(1,10)_ = 1.563, *p* = 0.2397) or change in the angle of slide (**Fig. 3E**, *F*_(1,10)_ = 0.9076, *p* = 0.3632) across cumulative ethanol doses. Assessment of righting reflex revealed no differences between AIR- and CIE-exposed mice (**Fig. 3F**, Pearson’s Chi Square, *p* = 0.2207). Thus, the ERB showed an increased physiological tolerance to ethanol in CIE mice only, evidenced by their blunted responsiveness to ethanol’s intoxicating and hypothermic effects despite having similar BECs to AIR-exposed mice.

**Figure 3.**
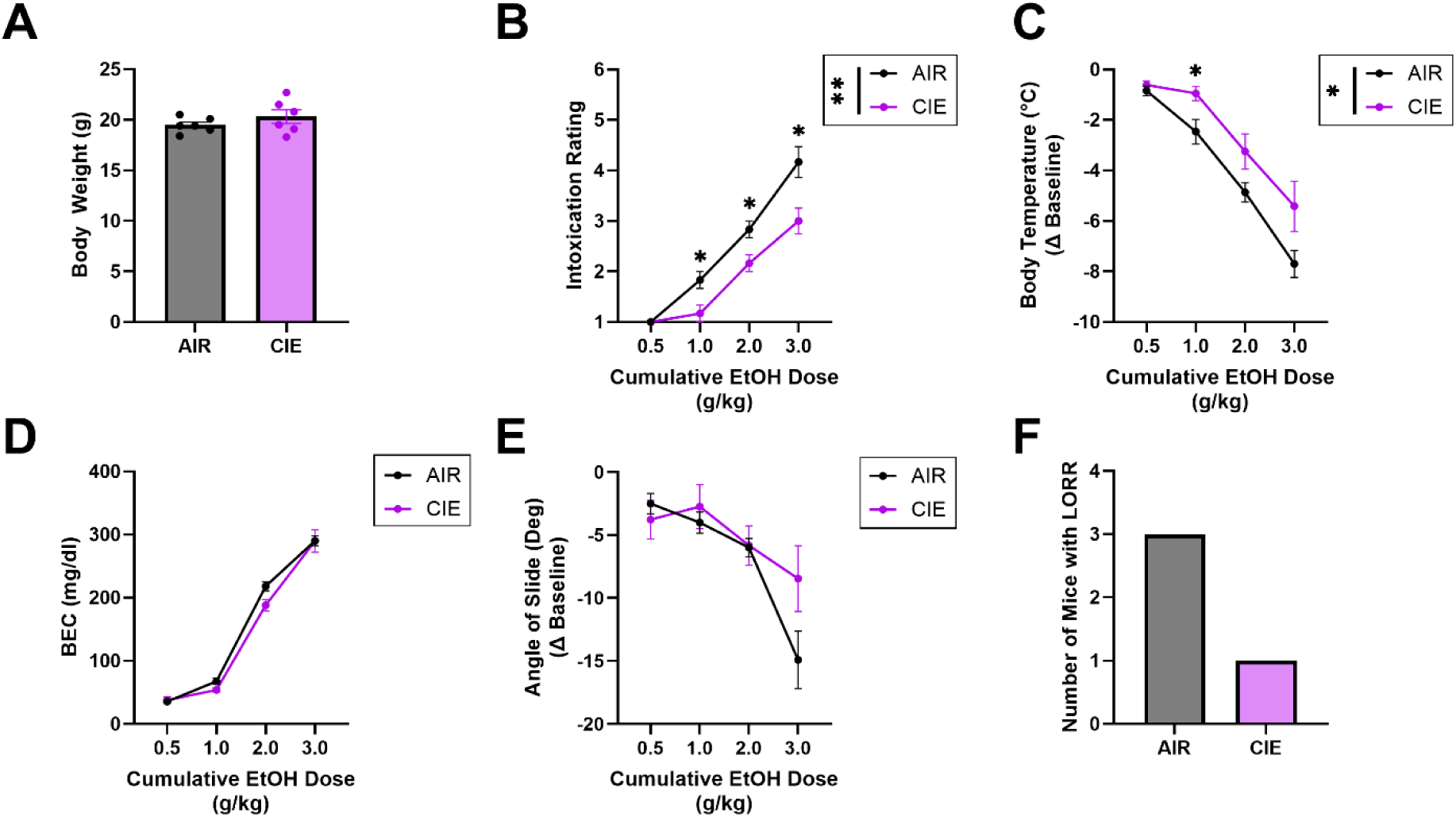
Effects of CIE exposure on ethanol responsivity **(A)** Body weights of 6 AIR- and 6 CIE-exposed mice one week after treatment, at the beginning of the ethanol responsivity battery (ERB). **(B-E)** ERB tests carried out after each intraperitoneal injection of ethanol, including **(B)** average intoxication ratings, on a scale from one to six, **(C)** change in rectal body temperature, **(D)** blood ethanol concentrations (BECs), and **(E)** change in the average angle at which each mouse would start sliding during the tilting plane assay. **(F)** Number of mice unable to right themselves during the loss of righting reflex (LORR) assay, which was only assessed after the final ethanol dose. Data are represented as mean ± SEM.

### CIE impact on elevated zero maze and open field

The effects of CIE on locomotion and anxiety-like behavior were assessed three and five days after completion of the CIE paradigm (**Fig. 4A**), using the elevated zero maze (EZM, **Fig. 4B**) and open field test (OFT, **Fig. 4F**) respectively. During the EZM assay, an unpaired t-test revealed a trend toward an increase in EZM open zone entries for CIE versus AIR mice (**Fig. 4C**, *t*(18) = 1.838, *p* = 0.0826). Additionally, the overall time spent in the closed zones was significantly greater than the time spent in the open zones (**Fig. 4D**, Two-Way RM ANOVA, F_(1,18)_ = 465.1, *p* < 0.0001), which was independent of group (*p* > 0.05). This difference was not due to locomotor effects, as the total distance traveled did not vary between groups (**Fig. 4E**, unpaired t-test, *t*(18) = 0.2175, *p* = 0.8303). For the OFT, CIE had no impact on the amount of time spent in the center of the open field (**Fig. 4G**, unpaired t-test, *t*(18) = 1.086, *p* = 0.2998), latency to enter the center (**Fig. 4H**, unpaired t-test, *t*(18) = 0.5452, *p* = 0.5923), or the total distance traveled during the 30-minute assay (**Fig. 4I**, unpaired t-test, *t*(18) = 1.571, *p* = 0.1337). However, a series of unpaired t-tests showed a significant effect of group on time spent exploring (*t*(18) = 2.387, *p* = 0.0282) and rearing (*t*(18) = 2.476, *p* = 0.0234), with CIE mice spending more time engaging in these behaviors than AIR mice. No difference was found in time spent grooming (unpaired t-test, *t*(18) = 0.4472, *p* = 0.6600). Overall, these tests reveal an effect of CIE abstinence on exploration, with CIE-exposed mice showcasing greater exploratory behaviors in the EZM and OFT assays.

**Figure 4.**
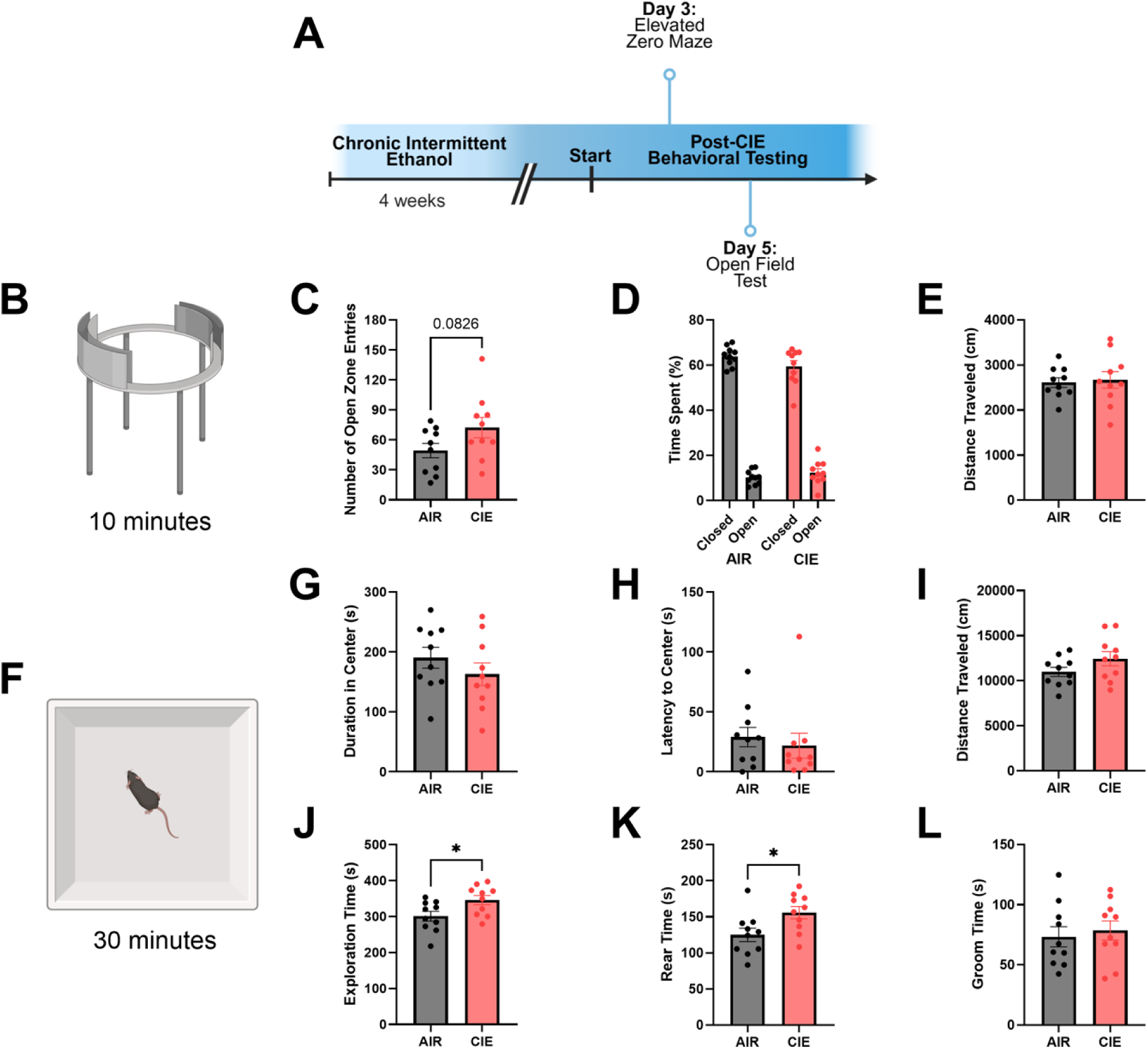
Effect of CIE on anxiety-like behaviors in the EZM and OFT. **(A)** Timeline of elevated zero maze (EZM) and open field test (OFT) behavioral assays after completion of the chronic intermittent ethanol (CIE) paradigm. **(B-E)** EZM assay, including **(B)** the set up. **(C)** Number of open zone entries throughout the duration of the assay. **(D)** Amount of time spent in the closed and open zones, calculated as a percentage of the total assay time. **(E)** Total distance traveled during the 10-minute assay. **(F-L)** OFT assay, including **(F)** the set up. **(G)** Amount of time spent in the center of the open field box. **(H)** Amount of time it took to enter the center of the open field. **(I)** Total distance traveled during the 30-minute assay. **(J-L)** Outcomes from ABEL behavioral scoring, including total time spent **(J)** exploring, **(K)** rearing, and **(L)** grooming. Data are represented as mean ± SEM.

### CIE leads to impaired novel object recognition in the NOR assay

To assess the impact of CIE on non-spatial memory, the novel object recognition (NOR, **Fig. 5B**) assay was conducted on Days 6 and 7 post-CIE (**Fig. 5A**). On the training day, a Two-Way RM ANOVA revealed no difference between AIR and CIE groups in the amount of time spent with either object (**Fig. 5C**, all *p* > 0.05). For the testing day, a Two-Way RM ANOVA revealed a significant effect of object (**Fig. 5D**, *F*_(1,18)_ = 11.24, *p* = 0.0035) and an interaction between treatment and object (*F*_(1,18)_ = 6.914, *p* = 0.0170). More specifically, Bonferroni’s *post hoc* test showed that the AIR mice spent more time with the novel object (*p* = 0.001), an effect that was not seen with the CIE mice. An unpaired t-test also showed that AIR mice had a significantly higher discrimination index than CIE mice on the testing day (**Fig. 5E**, *t*(18) = 2.395, *p* = 0.0277). In sum, this assay uncovered a significant blunting of the ability to discriminate between a novel and familiar object for CIE mice seven days after CIE-exposure.

**Figure 5.**
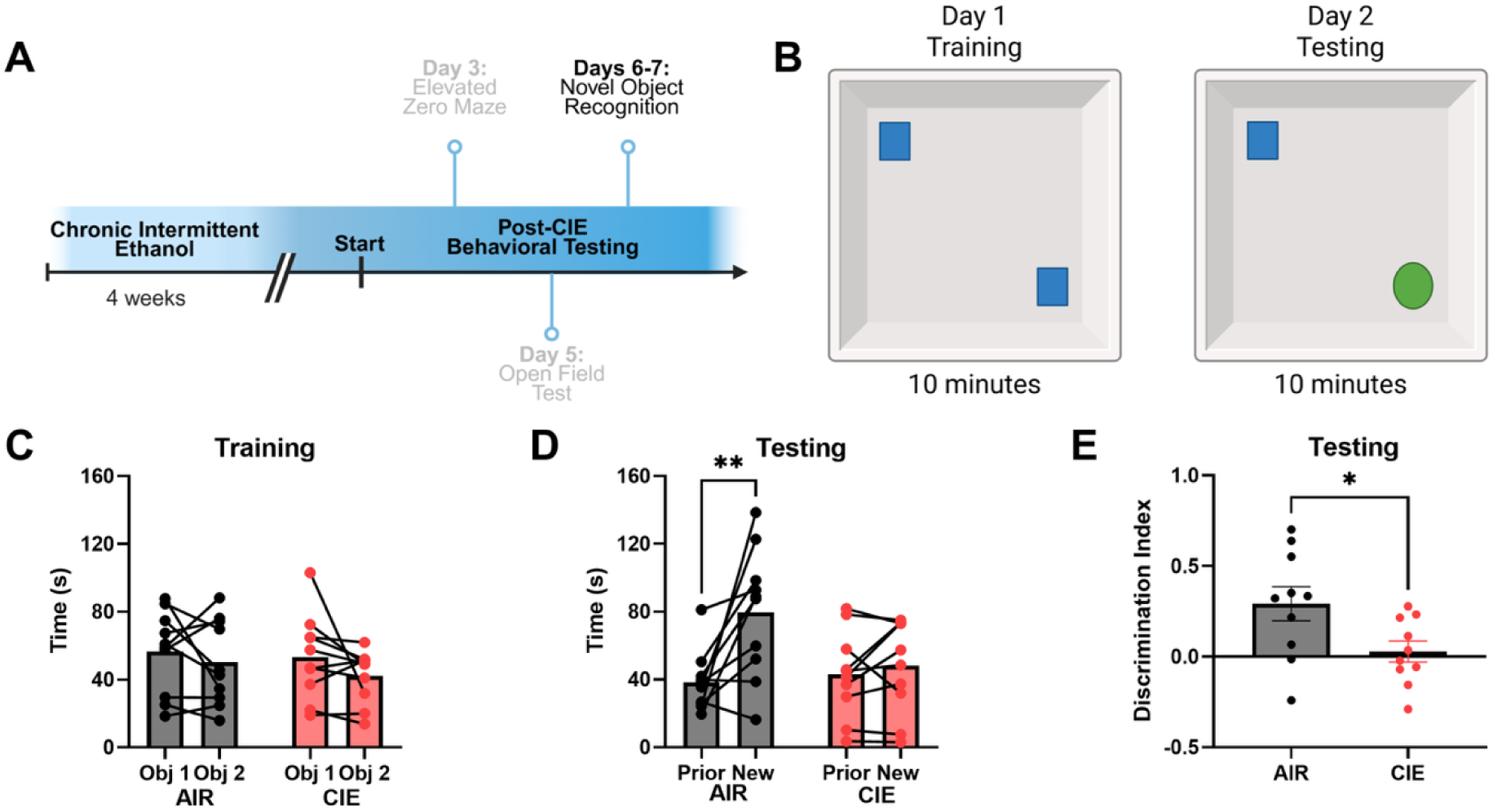
Effect of CIE-exposure on object recognition in the NOR assay. **(A)** Timeline of the novel object recognition (NOR) behavioral assay post-CIE. **(B)** Outline of the NOR experimental set up. Day 1 is also known as the training day where mice are given 10 minutes to explore two of the same objects. On day 2, the testing day, one of the objects from the training day is replaced with a novel object. The orientation of the objects remains the same between days. **(C)** Amount of time spent during the training day with object 1 versus object 2. **(D)** Amount of time spent during the testing day with the object from the training day, “prior,” versus the novel object, “new.” **(E)** The ability of the mice to discriminate between the “prior” and “new” object on the testing day. Data are represented as mean ± SEM.

### Impact of CIE on performance in light-dark box, novelty-induced suppression of feeding, acoustic startle and elevated plus maze

On post-CIE Days 12 through 21, anxiety-like behavior and startle responsivity of mice was assessed using the light-dark box (LDB, **Fig. 6B**), novelty suppressed feeding (NSF, **Fig. 6F**), acoustic startle (**Fig. 6H**), and elevated plus maze (EPM, **Fig. 6K**) assays. On Day 12 post-CIE, in the LDB assay, a series of unpaired t-tests showed no significant differences between CIE and AIR mice on the amount of time spent in the light side (**Fig. 6C**, *t*(18) = 0.5028, *p* = 0.6212), number of entries into the light side (**Fig. 6D**, *t*(18) = 0.3593, *p* = 0.7236), and latency to enter the dark side of the box (**Fig. 6E**, *t*(18) = 0.7153, *p* = 0.4836). For the NSF assay, ran on Day 14 post-CIE, CIE mice displayed a significantly greater latency to eat as compared to AIR mice (**Fig. 6G**, unpaired t-test, *t*(18) = 3.521, *p* = 0.0024). There was also no difference in appetitive drive between AIR and CIE mice, as they ate a similar amount of food (**Supplemental Fig. 1C**, unpaired t-test, *t*(18) = 0.7556, *p* = 0.4596).

**Figure 6.**
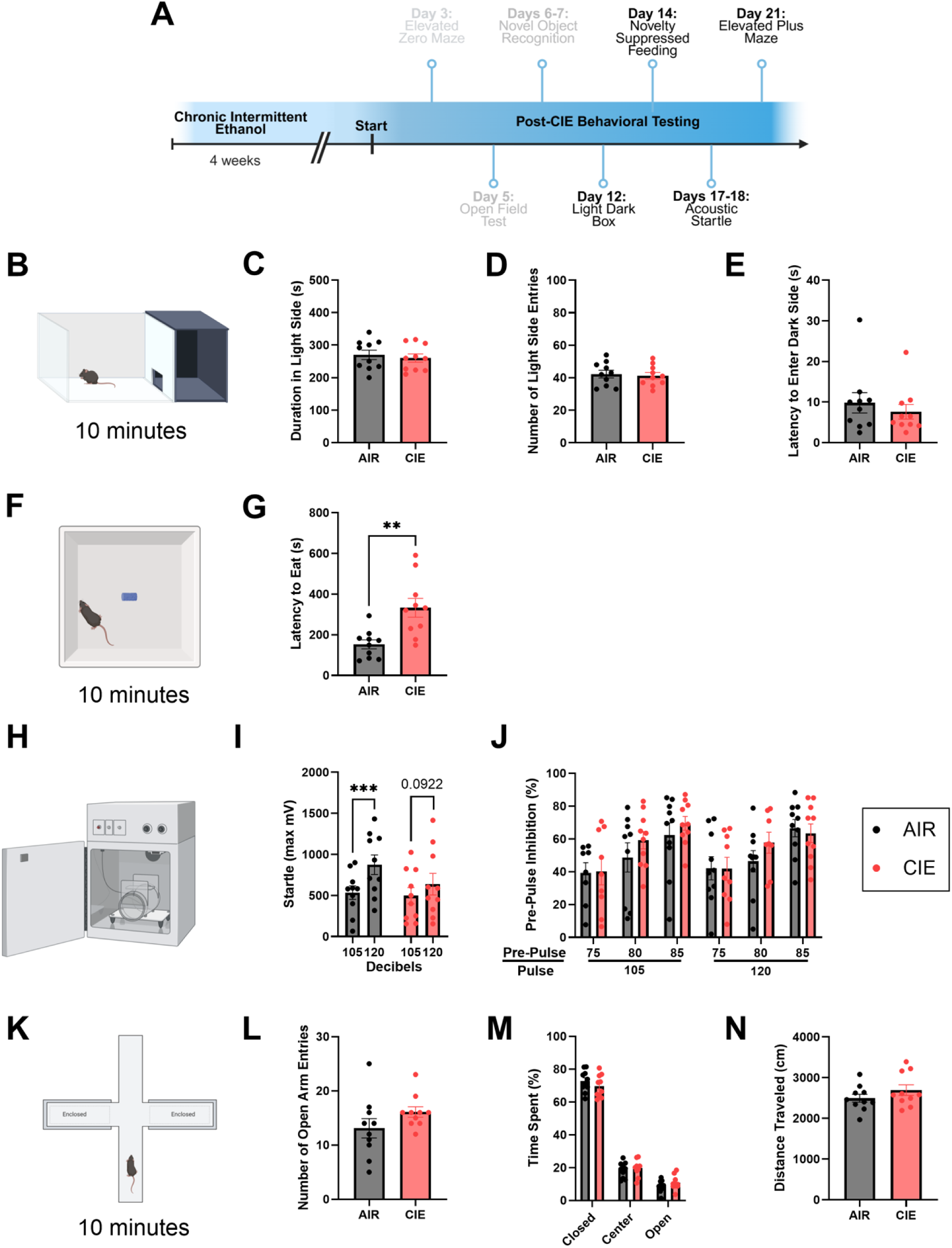
Effect of CIE-exposure on anxiety-like behavior in the LDB, NSF, Acoustic Startle, and EPM assays. **(A)** Timeline of light-dark box (LDB), novelty suppressed feeding (NSF), acoustic startle, and elevated plus maze (EPM) assays after completion of the chronic intermittent ethanol (CIE) paradigm. **(B-E)** LDB assay including **(B)** the set up. **(C)** Amount of time during the LDB assay that AIR and CIE mice spent in the light side. **(D)** Number of entries to the light side during the 10-minute assay. **(E)** Latency, or amount of time, to enter the dark side. **(F-H)** NSF assay including **(F)** the set up. **(G)** Amount of time for mice to eat the blue food pellet. **(H-J)** Acoustic startle assay including **(H)** the set up. **(I)** Mouse max startle responses to various decibel tones. **(J)** Pre-pulse inhibition for each pre-pulse/pulse pairing. **(K-N)** EPM assay including **(K)** the set up. **(L)** Frequency of entries into the open arm. **(M)** Amount of time spent in the closed arms, center, and open arms, calculated as a percentage of the total assay time. **(N)** Total distance traveled during the 10-minute assay. Data are represented as mean ± SEM.

In the acoustic startle assay, which was performed on Day 17 post-CIE, the AIR mice had a significant increase in startle response for the 120-decibel tone versus the 105-decibel tone, an effect that was not seen for the CIE mice (**Fig. 6I**). A Two-Way RM ANOVA showed a significant effect of decibel (*F*_(1,18)_ = 26.62, *p* < 0.0001) and a significant interaction between decibel and treatment group (*F*_(1,18)_ = 4.607, *p* = 0.0457). While there was a significant increase in pre-pulse inhibition across pre-pulses for both the 105- and 120-decibel tones (**Fig. 6J**, Two-Way RM ANOVA, *F*_(3.325,55.19)_ = 8.943, *p* < 0.0001), this increase was similar between AIR and CIE mice (Two-Way RM ANOVA, *F*_(1,18)_ = 0.7215, *p* = 0.4068).

On Day 21 post-CIE, for the EPM assay, unpaired t-tests revealed no differences in number of open arm entries (**Fig. 6L**, *t*(18) = 1.484, *p* = 0.1551) or locomotion (**Fig. 6N**, *t*(18) = 1.226, *p* = 0.2361) between treatment groups. A significant difference was found between the time spent in the closed, center, and open arms (**Fig. 6M**, Two-Way RM ANOVA, *F*_(1.424, 25.64)_ = 527.0, *p* < 0.0001), which was independent of treatment (Two-Way RM ANOVA, *p* > 0.05). Collectively, these set of assays reveal an increased anxiety-like effect of ethanol abstinence on post-CIE Day 14 in the NSF assay, and an impaired startle response to increasing tone decibels on post-CIE Days 17-18.

### Impact of CIE on cued fear conditioning and extinction

The effect of CIE-exposure on learning and memory was tested 25-27 days post-CIE (**Fig. 7A**) using cued fear conditioning (**Fig. 7B**). During the fear conditioning day, both AIR and CIE mice showed an increase in the amount of freezing across tone-shock presentations (**Fig. 7C**, Two-Way RM ANOVA, *F*_(5,90)_ = 44.70, *p* < 0.0001) which was independent of treatment (*F*_(1,18)_ = 0.7623, *p* = 0.3941). A Two-Way RM ANOVA also revealed an increase in freezing across tone presentations during the extinction day (**Fig. 7D**, *F*_(3.237,58.27)_ = 52.80, *p* < 0.0001) which was independent of treatment (*F*_(1,18)_ = 1.224, *p* = 0.2831). On the extinction recall day, there was a significant difference between the amount of freezing of AIR and CIE mice (**Fig. 7E**, Two-Way RM ANOVA, *F*_(1,18)_ = 4.547, *p* = 0.0470), which a Bonferroni’s *post hoc* test showed to be specifically on tone presentations 1-5 (*p* = 0.0102). A Two-Way RM ANOVA also revealed a significant effect of tone presentation on freezing (*F*_(2.707,48.73)_ = 21.44, *p* < 0.0001). In sum, this assay shows an effect of CIE on fear extinction recall, as CIE mice spent more time freezing than their AIR-exposed counterparts on extinction recall day.

**Figure 7.**
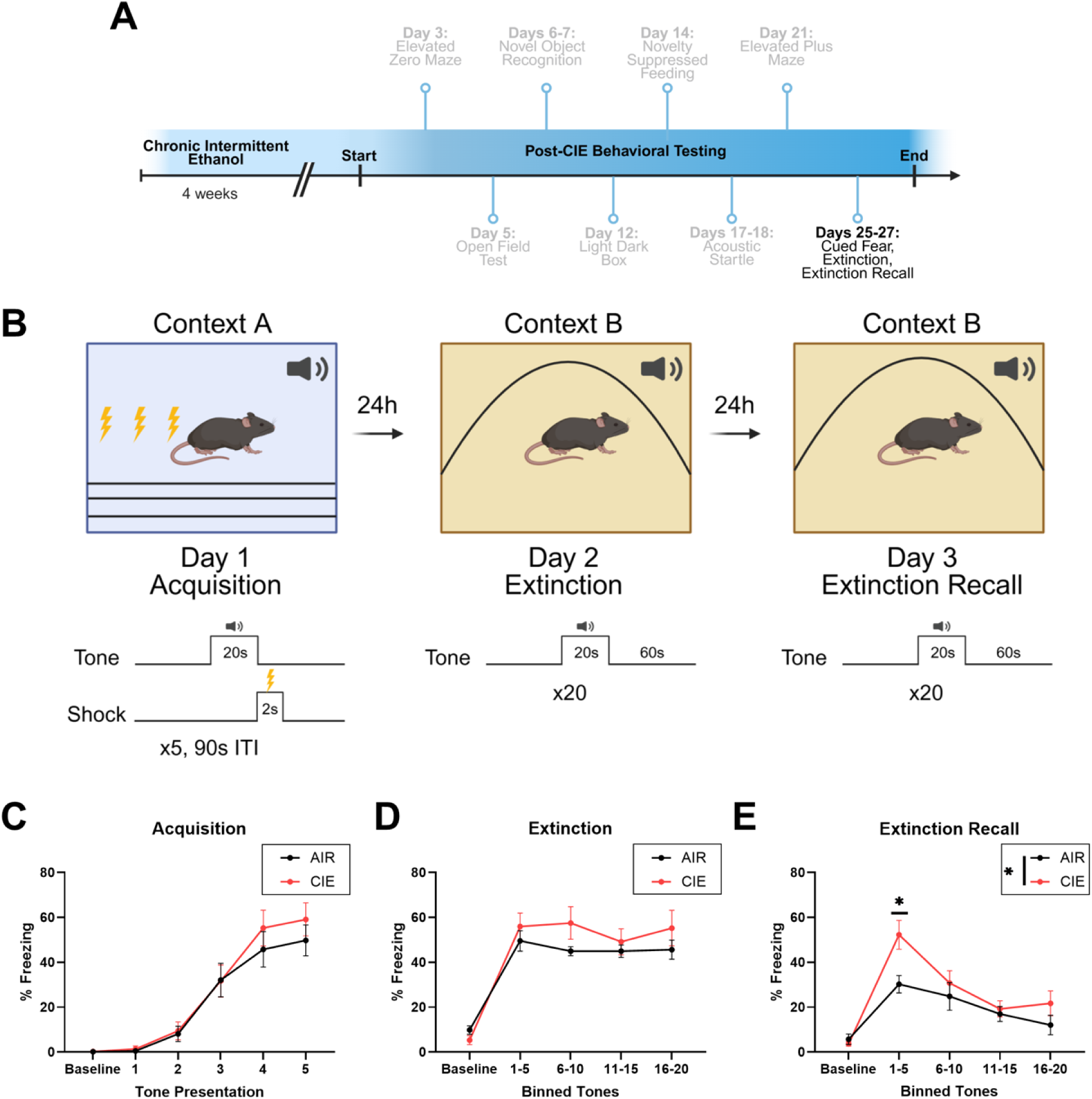
Impact of forced ethanol abstinence on cued fear conditioning **(A)** Timeline of cued fear conditioning during forced abstinence from the chronic intermittent ethanol (CIE) paradigm. **(B)** Model of fear conditioning protocol, which occurred in context A, as well as extinction and extinction recall, both of which were run in context B. **(C)** Percentage of time spent freezing across each tone presentation, during fear acquisition. **(D)** Average percentage of time spent freezing for every group of five tone presentations, during fear extinction. **(E)** Average percentage of time spent freezing for every five tone presentations, during extinction recall. Data are represented as mean ± SEM.

## Discussion

The present study used the chronic intermittent ethanol (CIE) vapor exposure paradigm to further understand the behavioral impact of ethanol abstinence in female mice. We evaluated both ethanol tolerance and changes in behavior, with an emphasis on more protracted time points, as these may provide critical translational insight.

In the first experiment, we focused on physiological tolerance to ethanol following CIE or air exposure, as there has been remarkably little exploration of this important phenotype. We saw a blunting of the sedating^37^ and hypothermic^38,39^ effects typically associated with chronic ethanol exposure, indicating an increased tolerance to the physiological effects of ethanol. This finding aligns with a previous study showing a reduction in loss of righting response in female mice exposed to CIE vapor^40^. However, it is notable that we observed this CIE-induced difference in tolerance at 7 days post-CIE, as opposed to 1 day post-CIE from Jury *et al.*^41^, suggesting that this change in tolerance persists beyond acute withdrawal. Moreover, we did not see any difference in BEC levels in CIE mice, indicating that this tolerance may be driven through central mechanisms rather than metabolic mechanisms. These results are important as heightened tolerance to ethanol is a key driver of excessive alcohol intake, and it supports the CIE model as an appropriate approach to study this phenotype.

In the second experiment, we wanted to evaluate changes in behavior following CIE-exposure during abstinence. As was highlighted in a recent review^42^, while there have been many studies exploring changes in affective behavior following alcohol exposure, there have been fewer investigations into how long these changes remain during abstinence, particularly with regard to the CIE model. Thus, to rigorously evaluate this, we evaluated changes in behavior across five weeks of abstinence following CIE-exposure. We focused our testing on probing anxiety-like behavior, as anxiety is a commonly reported symptom experienced during ethanol abstinence^43^, potentially due to the impact of ethanol on anxiety-related brain regions^44^.

Consistent with previous findings^19,21,45^, we saw no robust evidence of an anxiety-like state for female mice early in the abstinence period. However, it was notable that in a study exploring the impact of intermittent access to alcohol consumption, both male and female mice showed increase open arm entries 24 hours into forced abstinence^46^, similar to the trend we observed in the EZM. This suggests that this change may be reflective of a behavioral disinhibition. This particular phenotype may be short lived, as when we investigated performance in the OFT assay at Day 5 post-CIE, there was no difference in center time. However, at post-CIE Day 14, an anxiogenic effect of ethanol abstinence was revealed in the NSF assay, in which CIE-exposed mice displayed hyponeophagia. This behavioral change has been noted before in forced abstinence from ethanol, where ethanol-abstinent females displayed no anxiety-like behaviors in EPM, but then had hyponeophagia in the NSF assay at a later abstinence time point^47,48^. Importantly, our study showed that the anxiety-like phenotype did not persist in the EPM assay, which was administered to the mice one week after the NSF assay. Notably, this is in contrast to a study that found persistent changes in NSF 35 days into forced abstinence from chronic alcohol consumption^49^. Out of the three behavioral tests mentioned, the NSF assay is the only one to employ an approach-avoidance conflict^50^, suggesting that the inclusion of an appetitive component rather than only an innate avoidance-like component may be more sensitive to long term alcohol exposure.

In addition to exploring anxiety-like behavior, we wanted to evaluate metrics of cognitive performance, in particular memory. First, we used the NOR assay as a means to explore non-spatial long-term memory. We found that 7 days post-CIE, there was a significant reduction in the percent of time exploring the new object compared to a familiar object, suggestive of a disruption in long-term memory. This observation is consistent with findings from the Crews lab that adolescent alcohol exposure can disrupt memory processing in female rats^51^. However, this is not consistent with recent work showing that chronic ethanol consumption impairs NOR in male, but not female mSP rats^52^. Importantly, there are several notable differences including species, ethanol exposure, and abstinence which may contribute to the differing results. We also assessed cued fear conditioning and extinction as additional behavioral processes that are linked with memory and learning. We found no impact on cued fear conditioning nor extinction training; however, we saw a significant impairment of extinction recall at post-CIE Day 27. This builds on previous work in male mice that found that CIE-exposure led to impairments of fear extinction recall 4 days post-CIE^53^. Disruption of fear extinction recall has also been found following adolescent alcohol exposure ^54,55^, yet some work has found no effect in female rats following adolescent alcohol exposure^56^. Taken together, it suggests that male and female rodents may experience memory deficits with distinct temporal profiles, with female changes emerging during abstinence.

In this study, we found evidence for heightened tolerance following CIE-exposure in female mice, a time and assay dependent increase in anxiety-like behavior as measured via the NSF assay, and disrupted cognitive function in NOR and fear extinction recall. This is of particular interest, as the CIE protocol can lead to increased alcohol consumption. While a dominant theory in the field is that this escalation is due to increased negative affective states, our data suggest that CIE-exposure may engender this through multiple mechanisms, that are likely mediated via distinct neurochemical and anatomical substrates. The persistence of CIE-induced changes in behavior highlight the utility of the CIE model to probe the mechanisms that can drive these changes. One limitation of this work is that we only performed these experiments in female mice, which precludes an evaluation of sex-differences.

