## Supplemental Figure 1 for "Chronic Intermittent Ethanol Exposure Produces Physiological Tolerance and Distinct Behavioral Alterations Across Abstinence in Female Mice" (from another version of this preprint: version 1, 2026-10-05)

**Methods**

**Appetitive Drive Test**

On Day 24 post-CIE, all CIE and AIR mice were tested on their appetitive drive. After four hours of food deprivation, mice were single-housed in cages similar to their home cage and given 10 minutes to explore. Each new cage contained one single pellet of pre-weighed high fat diet (HFD, Research Diets D12492) that was placed on the floor of the cage. Once the session was over, the remaining HFD was weighed, and mice were returned to their home cage with their cage-mates.

**Figures**

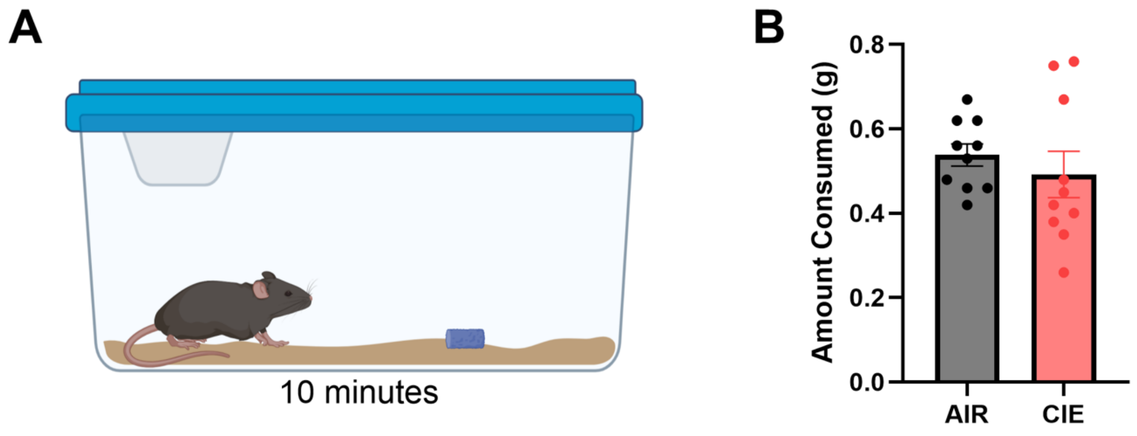


Supplemental Figure 1: Effect of CIE-exposure on the Appetitive Drive Test.

**(A)** Appetitive drive behavioral test set up, conducted 24 days post-CIE. **(B)** Amount of food consumed during the 10-minute appetitive drive test. Data are represented as mean ± SEM.
